# Distance and Colour-based Scores for Blood Test Risk Stratification

**DOI:** 10.1101/2020.02.09.941096

**Authors:** Hector Zenil, Francisco Hernández-Quiroz, Santiago Hérnandez-Orozco, Kourosh Saeb-Parsy, Héctor Hernández De la Cerda, Jürgen Riedel

**Affiliations:** Oxford Immune Algorithmics, Davidson House, The Forebury, RG1 3EU, U.K

## Abstract

We introduce a numerical and a colour-based risk stratification score to quantify abnormal blood analyte values. The score indicates how removed values of an individual are from considered healthy ranges from the literature or derived from empirical data such as medical surveys. The scores’ behaviour can be adjusted to incorporate medical knowledge by assigning multipliers or ‘weights’ to individual components and is rooted on a numerical and a colour-based scheme. We test the score against real and synthetic data from medically relevant cases, extremes cases, and empirical blood cell count data from the CDC NHANES survey spanning 13 years, from 2003 to 2016. We find that both the numerical and colour-based scores are informative in distinguishing healthy individuals from those with diseases manifested with abnormal blood results.

## 1 Background

An analyte or test parameter is a non-mutually exclusive property related to a blood test. Analytes can include protein-based substances, antibodies, biochemical entities and any other product or sub-product of cellular function related to the blood or to the immune system. For example, in a Full (or Complete) Blood Count, there will usually be about 13 to 15 analytes (see Table 1, although the score is not limited to any particular set or number of analytes. In the context of a cell’s properties, relevant tests can be, for example, cell count, cell size, cell morphology, cell nuclei morphology and cell maturity among a wide range of parameters/analytes.

**Table 1:** Most common analytes in a Complete or Full Blood Count with 5-part differential (CFC or FBC, hereafter simply FBC).

|  | **Analyte** | **Abbreviation** |
| --- | --- | --- |
| 1 | Haemoglobin | Hb |
| 2 | White Blood Cell count | WBC |
| 3 | Platelets count | Plt |
| 4 | Red Blood Cell count | RBC |
| 5 | Mean Cell Volume | MCV |
| 6 | Haematocrit | Hct |
| 7 | Mean Cell Haemoglobin | MCH |
| 8 | Mean Cell Haemoglobin Concentration | MCHC |
| 9 | Neutrophil count | N |
| 10 | Lymphocyte count | L |
| 11 | Monocyte count | M |
| 12 | Eosinophil count | E |
| 13 | Basophil count | B |

The complete blood count (CBC) with 5-part differential includes the number of red blood cells (RBCs), white blood cells (WBCs) and platelets; measures haemoglobin; estimates the red cells’ volume and sorts the WBCs into subtypes. A CBC is a routine blood test used to evaluate overall health and detect a wide range of disorders, including anaemia, infection and leukaemia.

The calculation of the score can incorporate normal ranges adjusted for age, race, gender, pregnancy stage, geography and any other consideration warranted by the literature.

### 1.1 Medical Considerations

The score aims to have clinical utility by capturing medical knowledge. In typical test results, normal reference values may be adapted for gender, age and pregnancy status. Rarely are they adapted by ethnicity and location, particular characteristics (e.g. smoker) or even temporal factors (e.g. time of day, which is known to introduce variations [4]).

The immune score has the option of incorporating a multiplier or ‘weight’ as a piece-wise function per analyte to modify its overall contribution relative to other analytes in a non-linear fashion. For example, in a Full Blood Count, conditions related to decreased white cell counts are milder than those associated with higher cell counts. However, the literature on conditions where there is a decrease of basophils, mast cells, monocytes and eosinophils in isolation, is sparse and reduced weights can be assigned to these markers.

Differences in cell shape can be clinically significant, but in the case of generally healthy individuals they are not given greater weight than differences in cell count. Typically, disorders affecting bone marrow function (i.e., blood cancers) result in the presence of abnormal (often immature) cells in peripheral blood; their presence in peripheral blood beyond this level would almost always be abnormal. A related point is whether the score reflects subtle/minor changes in shape (variation in cell size, nuclear size, presence of other organelles) or only very crude and major deviations in size. For white blood cells, we do not yet understand what significance these have.

The score is not intended to be used as a diagnostic tool and can only quantify abnormality by deviation from healthy reference values. The score is sensitive to out-of-range values and increases its value or changes its colour as a function of how removed values are from lower and upper bounds healthy reference values according to the number of standard deviations from the medians, but cannot quantify diseases or conditions. The score indicates how far the bulk of all markers are from normal (healthy) reference values, the median, and an interval determined by published reference values for specific demographic or health conditions. Medians can be derived from empirical data (see Fig 7 in the Complementary Material).

### 1.2 General description and notation

The immune score constitutes a dimensional reduction technique based on a single real-value number and a colour scheme that takes a multidimensional blood test space with dimension size equal to the number of analytes, where each value of a marker or analyte corresponds to the coordinate of that value in that dimension. The score itself can be seen as the norm of a suitable transformation of a vector pinpointing the health status of a patient in that space, relative to that test and set of markers or analytes. So the numerical vector value integrates all the analyte space dimensions.

Let

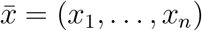

be a vector of raw analyte values obtained by a suitable (series of) lab tests, as usually presented to physicians and patients (see Table 1).

We call the associated vector space for blood cell analytes the *multidimensional immune space*, or the *immune space* for short.

## 2 Methods

The CDC NHANES data was used to estimate deficiencies and toxicities of specific nutrients in the population and subgroups in the U.S., to collect population reference data and to estimate the contribution of diet, supplements and other factors to whole blood levels of nutrients. Data can be used for research purposes and is publicly available (see e.g. Fig. 7 in the Complementary Material).

### 2.1 Numerical score

This first description of the score starts as a linear function because it does not take into account possible interactions between analytes. We also assume that each analyte contributes equally to the immune score (this is likely only partially true, as some of the analytes are more or less medically informative than others, with this informativeness itself varying with different conditions. Moreover, not all analytes are independent and some analytes may be statistically dependent on others—suitable refinements will be introduced later).

For ease of comparison between successive versions of the score, all values are normalised within the range [0, 10]. Which means that each analyte will contribute with a maximum weight of

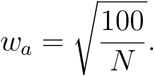

where *N* is the number of analytes. From now on we will consider the example of *N* = 13 typical for a Full Blood Count (FBC) test. From a geometrical point of view, a vector of values in the multidimensional immune space will be mapped to a point in a *N* D-“quadrant” going from 0 to *w*_*a*_ that will be called the *normalised immune space*. This view will allow us to observe the diachronic evolution of an individual’s score as a trajectory in the normalised immune space where a notion of distance will come in handy.

The adoption of a score within a maximum range of [0, 10] can be done in different ways. One approach is to assume no theoretical maximum values for analytes and asymptotically approach the maximum score of 10, without ever reaching it in individual cases. We adopted an alternative approach, capping values to a pre-established “reasonable” maximum beyond which all specific values mean the same. The pre-established maximum is two standard deviations beyond normal (healthy) ranges.

In an initial test, reference tables from *Hematology Reference Ranges* from the NHS (see Table 5 in the Complementary Material) were used for normal (healthy) ranges for each analyte according to age, sex and pregnancy status [3]. *G* will denote the set of possible categories:

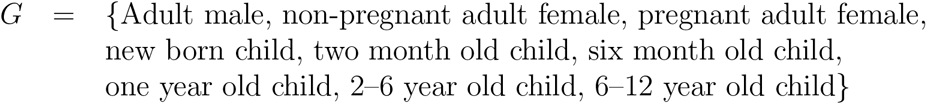

Let *g* denote any value in *G*.

For each analyte, *r*_*u*_(*g, i*) is the upper limit in the normal range of the analyte *i* for an individual belonging to group *g*. In a similar way, *r*_*l*_(*g, i*) is the lower limit.

Pre-calculate the following values:

- The *expected vector*, containing the mean value of each analyte for a given group:

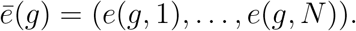

Mean values will come from the analysis of suitable data to be obtained from reliable databases or directly collected by us. In the meantime, we are taking the arithmetical average of the lower and upper limits in the NSH table:

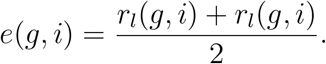
- The standard deviation for an analyte *i* in group *g* is denoted by

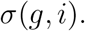

Again, these values will come from a suitable set of cases. In the meantime, we will use the distance from the mean to either limit within the normal (healthy) range:

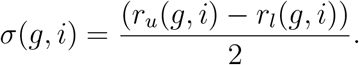
- The *maximum difference vector*, containing the maximum possible distance from the mean value of each analyte:

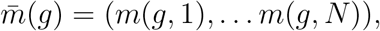

where

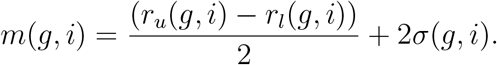

(twice standard deviations from the limits of the healthy ranges).
- The *weighted vectors* of each group (which will be used to normalise the possible values to a maximum immune score of 10):

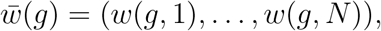

where *w*(*g, i*) = *w_a_/m*(*g, i*).

Let *c̄* = (*c*_1_,…,*c*_*N*_) be a vector of analyte values for an individual of group *g*. Then the normalised vector is calculated using the following formula

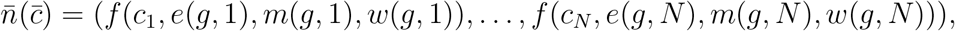

where

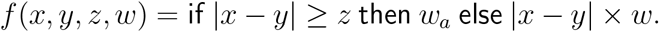

The norm of the normalised vector will be called the Normalised Immune Score, or NIS for short:

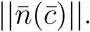

As shown in Fig. 1, the behaviour of the numerical value is consistent through varying number of analytes and scales linearly with respect of the deviation from the norm. This 2-D rectilinear manifold space where immune score values can lay. Fig. 1 explores the behaviour of the immune score over synthetically generated patient values as a function of number of analytes and removal from healthy reference values.

**Figure 1:**
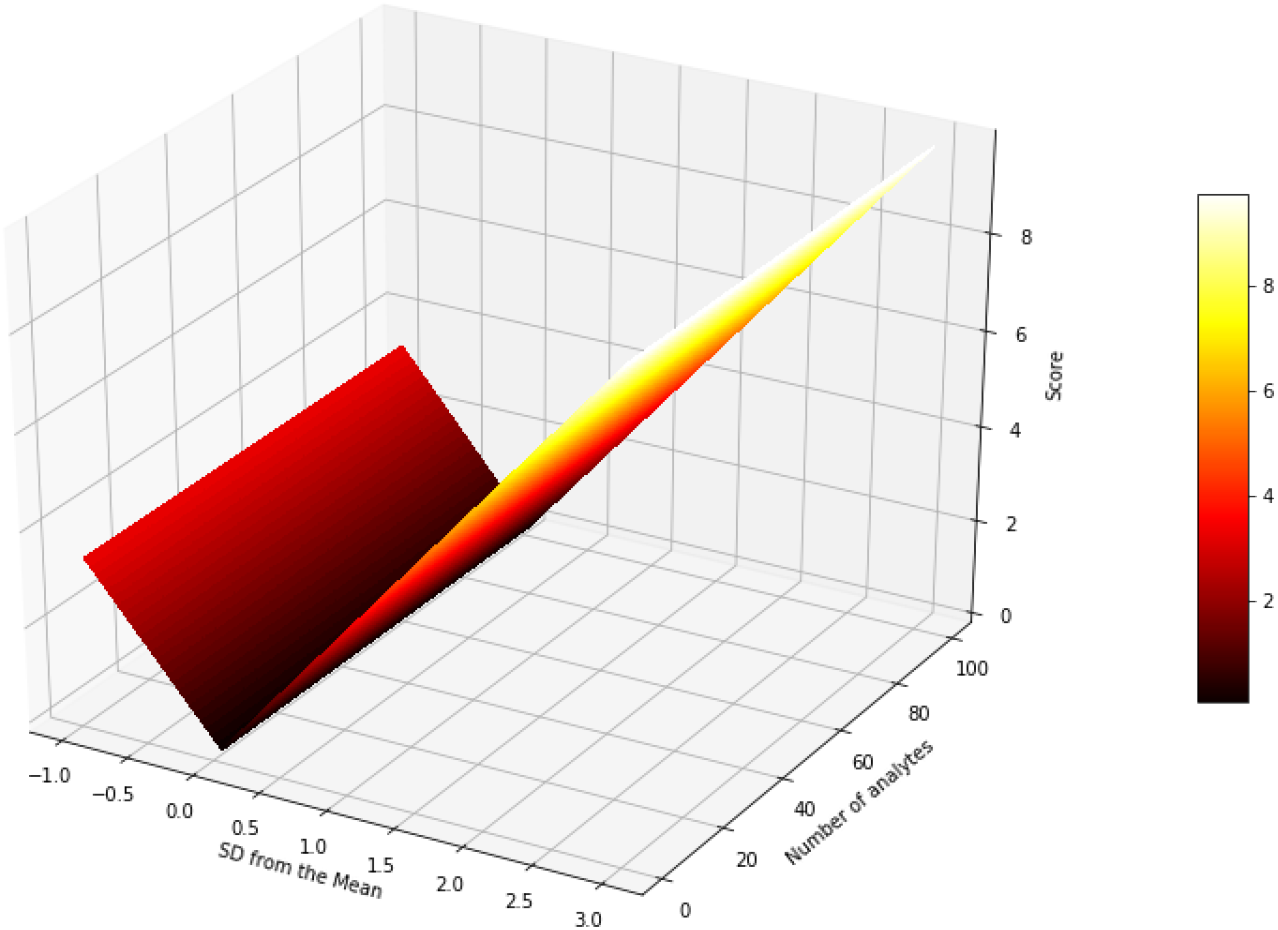
Immune score space: The *x* axis represents the deviation of all the analytes from the expected value, while the *y* axis measures the total number of analytes. A *negative* standard deviation indicates that the value is lower than expected and a positive one indicates a larger value than expected. The cut-off seen in the negative *x* axis shows the fact that real life values for analytes have an strict lower bound but, technically, have no upper bounds. For instance, is not possible to have a negative white blood cell count.

### 2.2 Colour-based score

In addition to this numerical value, the score incorporates a second indicator that will be easy to read and interpret. The concept is to utilise colour codes as a means of flagging deviations from normal (healthy) reference values. This is called the *Colour-Coded Immune Score* or CCIS for short.

The CCIS uses the same analytes and their normal (healthy) ranges as before. There are thresholds for each of the analytes that trigger different types of alerts:

- A value within normal healthy ranges: No alert, colour coded *green*.
- An abnormal value within one standard deviation above/below healthy reference values: *amber* alert.
- An abnormal value beyond one standard deviation of the normal healthy values: *red* alert. As in the normalised immune score, values are capped to a maximum of two standard deviations.

The results of each analyte are also combined in a single global value, which will constitute the CCIS. The rules are as follows:

- Green: all values within normal range (green).
- Amber: 1–3 amber individual alerts.
- Red: More than 3 amber alerts or one or more red alerts.

As can be seen, the CCIS is not a numerical value, but a colour-coded output, like the individual alerts for separate analytes. This clearly distinguishes the CCIS from the normalised immune score.

The information for the CCIS can be presented in a simple graphical way. We construct a doughnut graph with one slice for each analyte. Analytes receiving the same colour will be grouped together, which will produce doughnut graphs of at most 3 different colour-coded sections.

Some examples for illustration are presented in Figs. 2–3 (without the corresponding numerical values):

**Figure 2:**
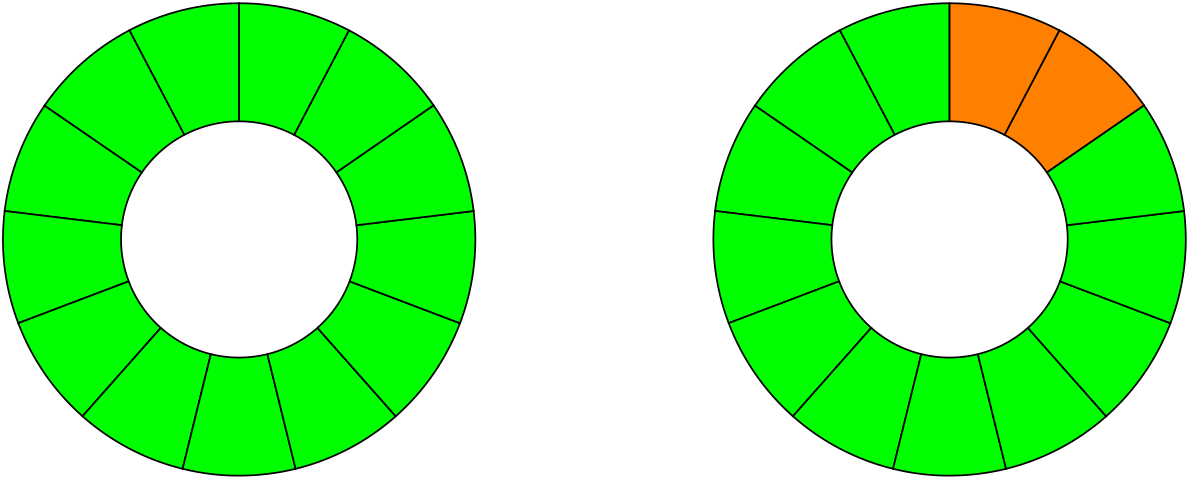
CCIS colour-based alerts. On the left, an individual with all values normal (healthy) and thus coloured green. On the right hand side, an individual with two amber alerts.

**Figure 3:**
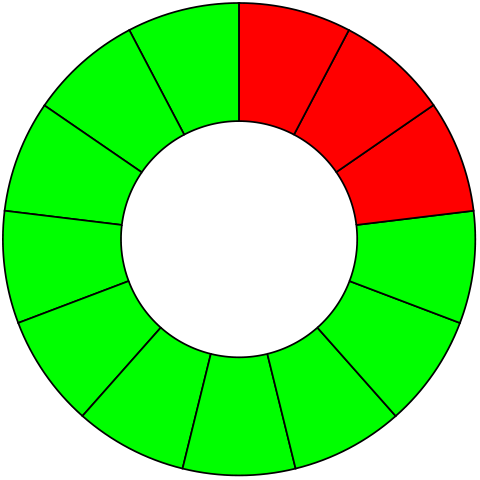
As in Fig. 2, on the left hand side there is a doughnut plot with three red alerts which may produce a global red flag quantifying potential risk, by colouring the numerical score that can be depicted in the centre of this colour-based score as a non-linear property.

## 3 A numerical score derived from the CCIS

We explored an alternative score that would follow the colour-coded scheme, but would also produce a numerical value. The intention was to combine the accessibility of the CCIS and the finer precision of the NIS in a single value. As with the latter, the value would range from 0 to 10.

We expected the following property: if *v*_1_*, v*_2_ are possible score values and *v*_1_ < *v*_2_, it should be the case that *v*_2_ indicates greater cause for concern than *v*_1_.

On the other hand, values should be closely related to the colour codes. That is, values *v*_1_, *v*_2_ and *v*_3_ correspond to CCIS scores of green, amber and red, respectively, and they should be ordered as follows:

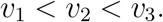

A more precise way of expressing the above ideas is the set of rules codified in Table 2.

**Table 2:** Mapping of score segments and colours.

| Colour from CCIS | Ideal interval for a numerical score |
| --- | --- |
| Green | $[1, 10/3]$ |
| Amber | $(10/3, 20/3]$ |
| Red | $(20/3, 10]$ |

In the following section, we present a procedure for translating the CCIS into a numerical value according to the above scenario. We will call this value the NCCIS. The expected behaviour of NCCIS with respect to the number of analytes and its deviation from the norm can be observed in Fig. 4.

**Figure 4:**
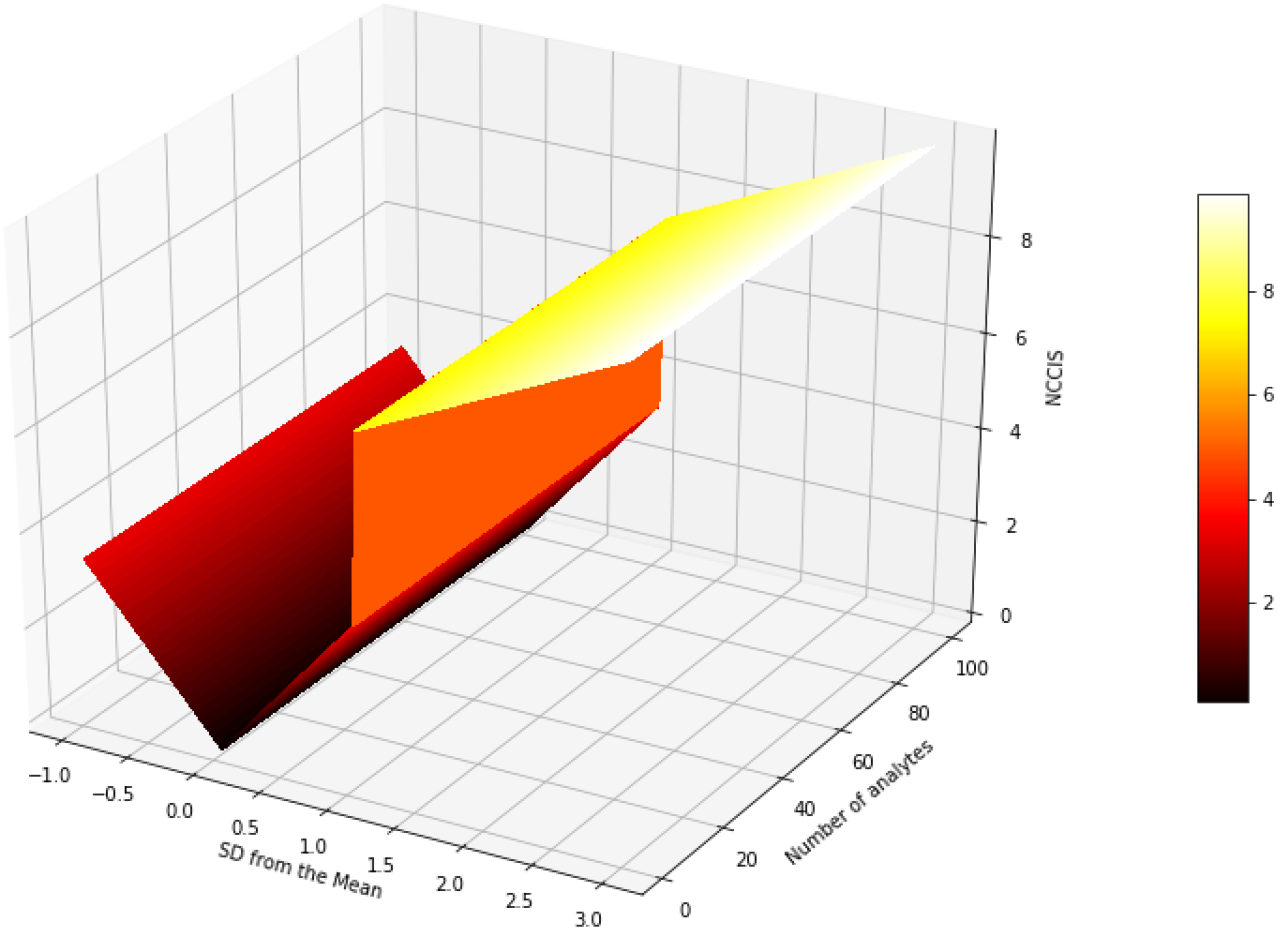
The behaviour of NCCIS over synthetically generated patients with differing values and a deviation from the norm with respect of the respective expected value.The *x* axis measures the stated while the *y* axis measures the total number of analytes. In contrast to the behaviour of the immune score (Fig. 1), NCCISS is a non-linear function that displays an *step-like* behaviour by design, while the independence towards the number of analytes remains.

The behaviour of Fig. 4, illustrates how values removed from healthy values are pushed towards higher values by design as an alerting mechanism.

### 3.1 Calculating CCIS and NCCIS

Some of the values in the calculation of the NIS:

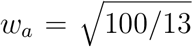 is the maximum normalised distance for each analyte in the normalised vector of a patient’s readings.
*e*(*g, i*) is the mean (or expected) value for analyte *i* in group *g*.
*σ*(*g, i*) is the standard deviation from the mean.
*n*(*g, i*) = (*r*_*u*_(*g, i*) – *r*_*l*_(*g, i*))/2 is the normal (healthy reference) interval length from the mean value (assuming upper and lower limits are equidistant from the mean, an assumption to be revised in the future both in the NIS and here in the NCCIS).

Now the raw (not normalised) borders between colours are

raw maximum value for green: *n*(*g, i*).
raw maximum value for amber: *n*(*g, i*) + *σ*(*g, i*).
raw maximum value for red: *n*(*g, i*) + 2*σ*(*g, i*).

And the normalised version:

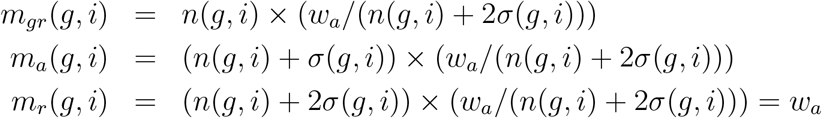

Therefore the current NIS intervals for the different colours for individual analytes are:

- Green: [0*, m_gr_*(*g, i*)] (that is, from 0 distance from the mean value up to the normalised maximum value for green.)
- Amber: (*m*_*gr*_(*g, i*)*, m_a_*(*g, i*)].
- Red: (*m*_*r*_(*g, i*)*, w_a_*].

For global values the calculation has to take into account both the definition of the CCIS and the fact that each analyte can have proportionally different normal ranges and standard deviations:

- *Green*: The minimum global green corresponds to 0 in every analyte. The maximum is *m*_*gr*_(*g, i*) in each analyte. This gives us the following interval:

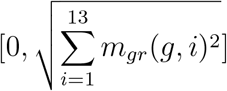
- *Amber*: The minimum global amber is 1 analyte value, just above green. The maximum is 3 maximum amber values and the rest (10) at the upper limit of green. Let analyte *j* be such that *m*_*gr*_(*g, j*) is the minimum of the green upper limits, and let analytes *k*, *m*, *n* be the three biggest of the amber upper limits. Then we have the following interval:

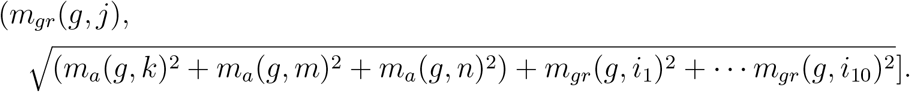
- *Red*. The minimum global red is 1 red analyte value or 3 amber analyte values. The maximum is, obviously, 10 (13 maximum individual analyte values). Let us suppose that the lowest minimum for red is (*m*_*r*_(*g, k*)) and that there are no 3 upper limits for amber analytes whose sum is below this. Then the intervals are:

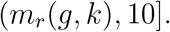

We will call the ends of these intervals min_*green*_, max_*green*_, min_*amber*_, max_*amber*_, min_*red*_ and max_*red*_.

The NCCIS is calculated using the following function:

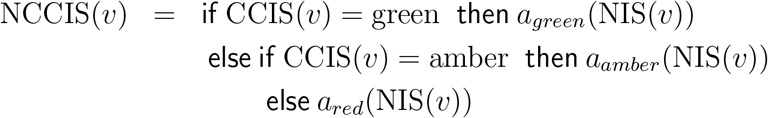

where CCIS and NIS are functions calculating the respective scores and the functions *g*_*a*_, *a*_*a*_ and *r*_*a*_ map the NIS to the intervals set in the table at the beginning of section 4:

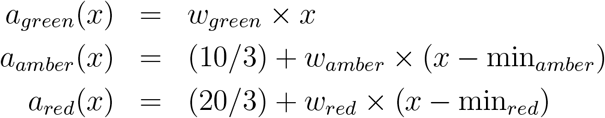

and the NIS value is weighted according to a normalising weight:

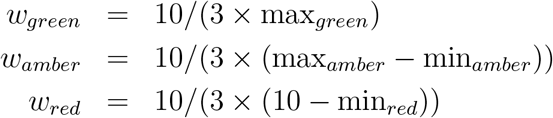

## 4 Numerical tests

In Table 3, we give CCIS and its correspondent NCCIS values for a sample of real life and artificial cases. The second column of the table also gives the NIS, for purposes of comparison. It is worth noting that that these examples were generated using the NHS values for normal healthy values (see 5 Complementary Material), and based on the provisional assumption that the distance between the upper and normal ranges is twice the standard deviation. The examples were taken from [2] as they were intended as very preliminary tests. We are aware that their source clearly states they are meant only for teaching purposes. Here they are used for purposes of illustration only and the development of the NIS and NCCIS is not dependent on them.

**Table 3:**
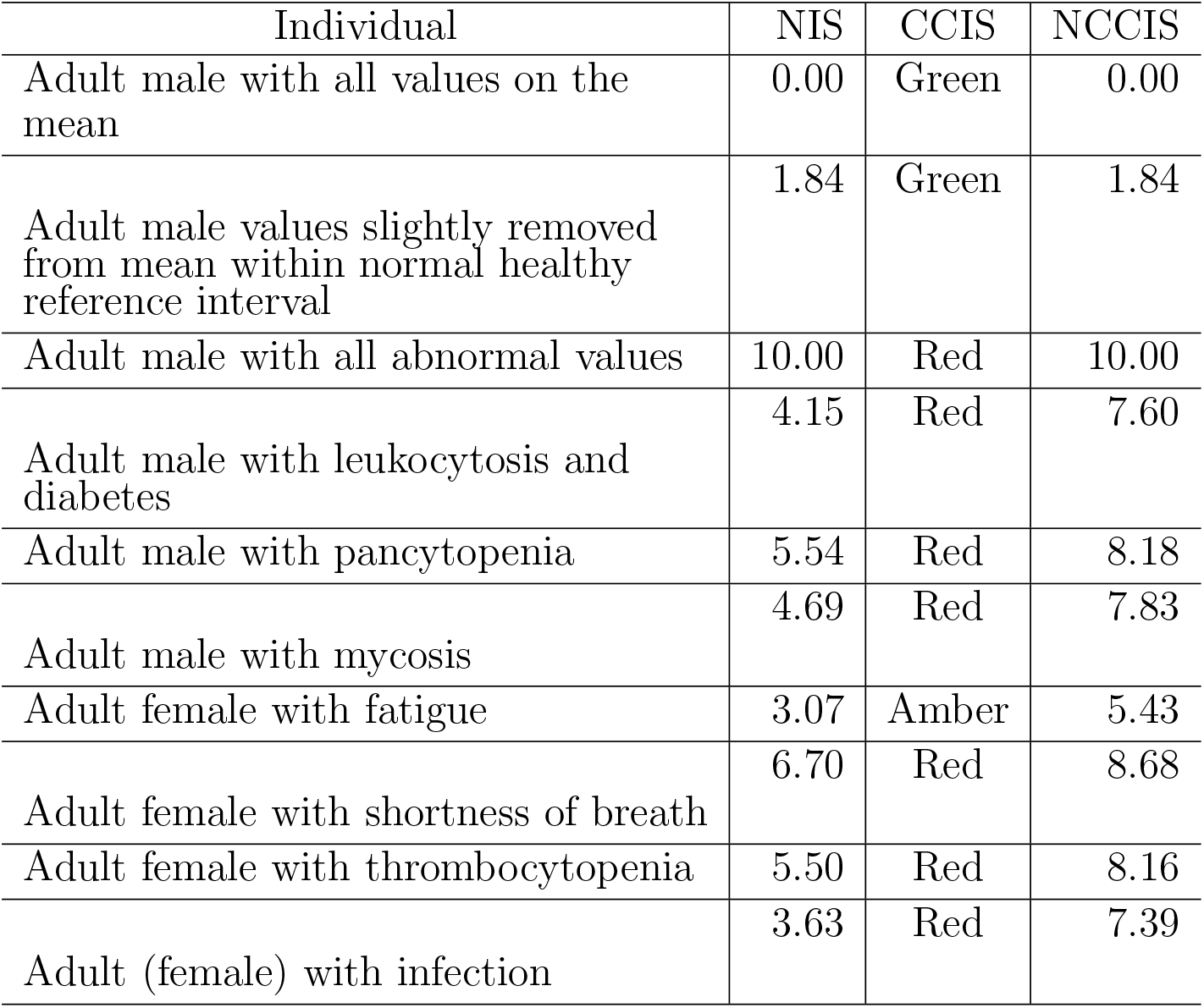
Different tested scores performing differently according to each score’s definition.

One can observe how the NCCIS values meet the requirements we set in advance of its definition. We tested the NIS and NCCIS against real data from the National Health and Nutrition Examination Survey 2003–2016 (NHANES), provided in [1]. One measure of success was their ability to show how accurately the score discriminates between healthy subjects and patients suffering from various diagnosed illnesses.

For this, we examined thousands of cases from [1] and calculated the distribution, mean values and standard deviations for each of the analytes (see Fig. 7 in the Complementary Material). We used the means and standard deviations from the data to define the metrics for both the NIS and NCCIS. Finally, we calculated the NIS and NCCIS for thousands of cases in the NHANES database to find individual scores. The resulting values were grouped as belonging to healthy individuals or to a selected list of common diseases.

In order to maximise the discriminatory power of both scores, we tried different alternative combinations of means, normal ranges and standard deviations, either taken from NHS tables (Table 5) (or inferred from them in the case of standard deviations e.g. Table 4 and Fig. 7 in the Complementary Material) or calculated from the NHANES database. At the end of the day, we settled for (1) NHS normal healthy reference values and means and standard deviations for the NIS; (2) means and standard deviations calculated from the NHANES database, and normal ranges from the NHS for the NCCIS. NHANES means and standard deviations are shown in table 4.

**Table 4:** Mean values and standard deviations found in the NHANES 2003– 2016 database for the 13 analytes of the immune scores.

| Analyte | Mean Male | Std Male | Mean Female | Std Female |
| --- | --- | --- | --- | --- |
| Hb | 150.72 | 11.51 | 133.33 | 11.62 |
| WBC | 6.92 | 3.12 | 7.01 | 2.09 |
| Plt | 228 | 54.33 | 255.42 | 64.19 |
| RBC | 4.93 | 0.45 | 4.40 | 0.38 |
| MCV | 89.96 | 5.13 | 89.22 | 5.72 |
| Hct | 0.44 | 0.03 | 0.39 | 0.03 |
| MCH | 26.96 | 10.33 | 26.55 | 10.47 |
| MCHC | 336.55 | 18.41 | 335.02 | 20.46 |
| N | 4 | 1.85 | 4.16 | 1.66 |
| L | 2.11 | 2.21 | 2.11 | 0.79 |
| M | 0.57 | 0.21 | 0.52 | 0.18 |
| E | 0.20 | 0.16 | 0.17 | 0.14 |
| B | 0.04 | 0.06 | 0.04 | 0.05 |

**Table 5:** NHS reference values: Typical tabular presentation of a Complete/Full Blood Count test.

| Parameter/Component | Result | Low Ref Value | High Ref Value |
| --- | --- | --- | --- |
| Haemoglobin | 152.3 | 115 | 165 |
| Total White Cell Count | 9.3 | 3.6 | 11.00 |
| Platelet count | 286.5 | 140 | 400 |
| Neutrophils | 6.9 | 1.8 | 7.5 |
| Lymphocytes | 3.6 | 1.0 | 4.0 |
| Monocytes | 0.3 | 0.2 | 0.8 |
| Eosinophils | 0.2 | 0.1 | 0.4 |
| Basophils | 0.2 | 0.02 | 0.1 |
| Red cell count | 5.5 | 3.8 | 5.8 |
| Haematocrit | 0.4 | 0.37 | 0.47 |
| Mean Cell Volume (MCV) | 93.3 | 80 | 100 |

In Fig. 5 we show how NIS values are distributed according to different (self-declared) conditions for individuals in the NHANES database. It must be noted that the self-reported conditions were not independently confirmed. Moreover, the survey participants did not distinguish between current or past diagnoses. This would be expected to introduce some noise, as some currently healthy people will be labelled with a condition and some people with conditions will not have been diagnosed. We hope to improve accuracy by filtering the data or by adding some other data sources in the future.

**Figure 5:**
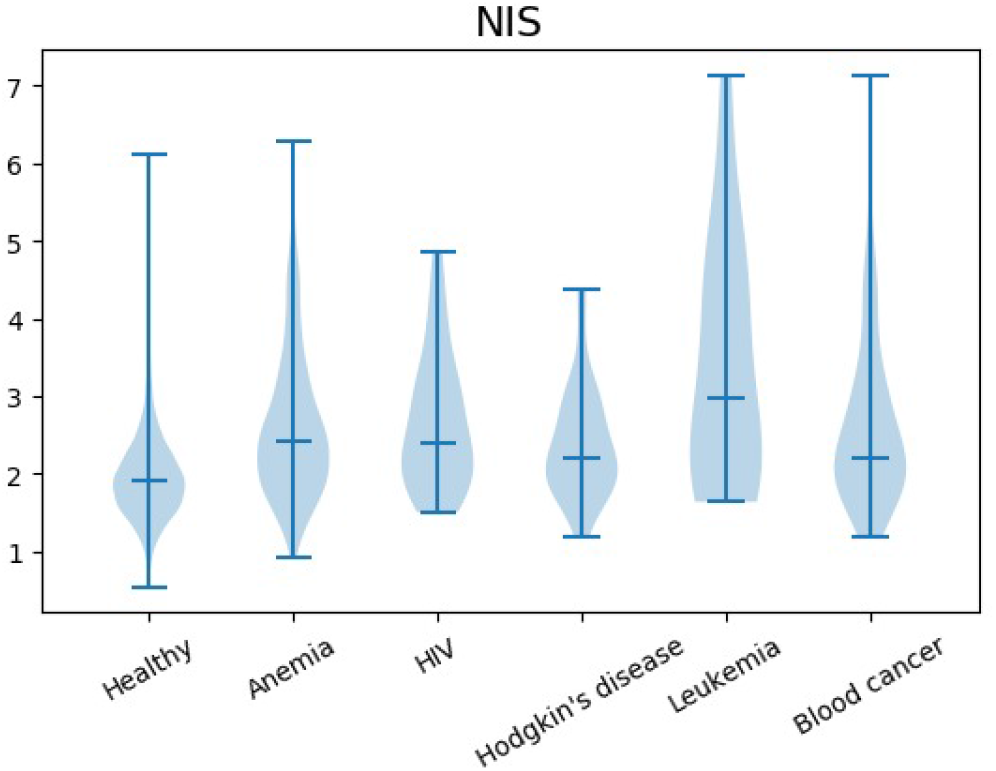
Distribution of the Normalised Immune Score (NIS) from 0 to 10 (*y* axis) among individuals in the NHANES 2003–2016 database.

Nevertheless, the NIS was able to discriminate between healthy and unhealthy individuals, as most healthy individuals are clustered around very low NIS values. In contrast, different conditions produced higher NIS values on average.

Conversely, the NCCIS did not substantially improve our knowledge of how some diseases impact analyte counts. In Fig. 6, it can be seen that medical conditions did produce a higher average NCCIS value, but also induced some anomalous clustering. Furthermore, the distribution of scores were even wider than that of the NIS. We thus concluded that the NCCIS did advance the knowledge already gleaned from the NIS and the CCIS.

**Figure 6:**
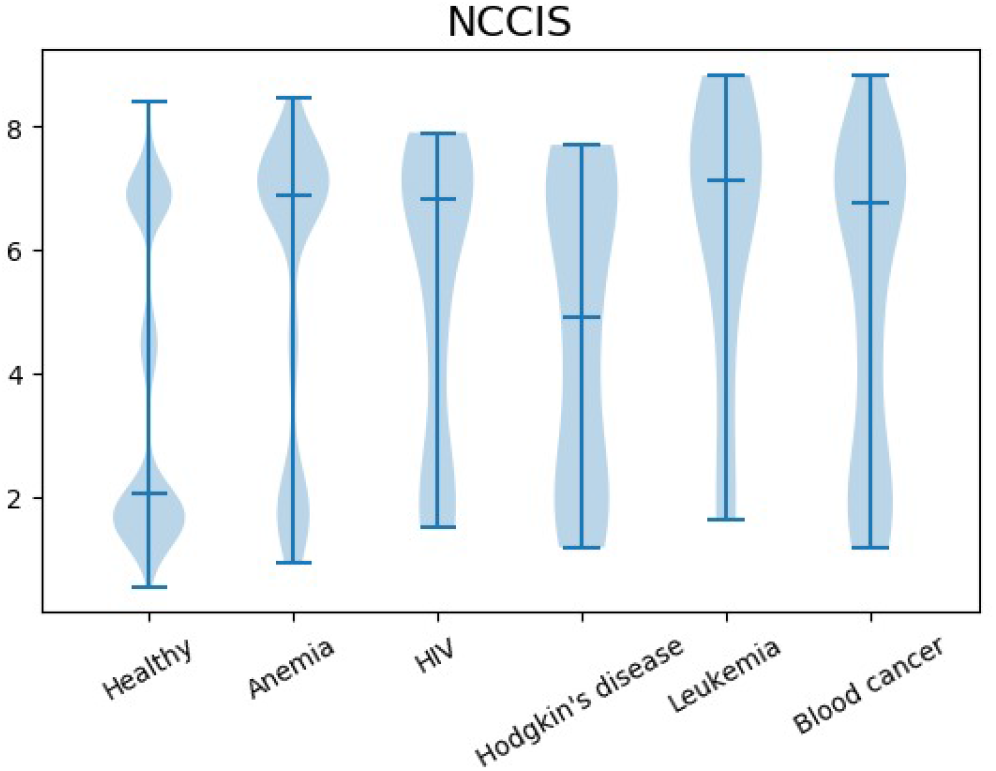
Distribution of the Normalised Colour-Coded Immune Score (NCCIS) from 0 to 10 (*y* axis) among individuals in the NHANES 2003–2016 database.

**Figure 7:**
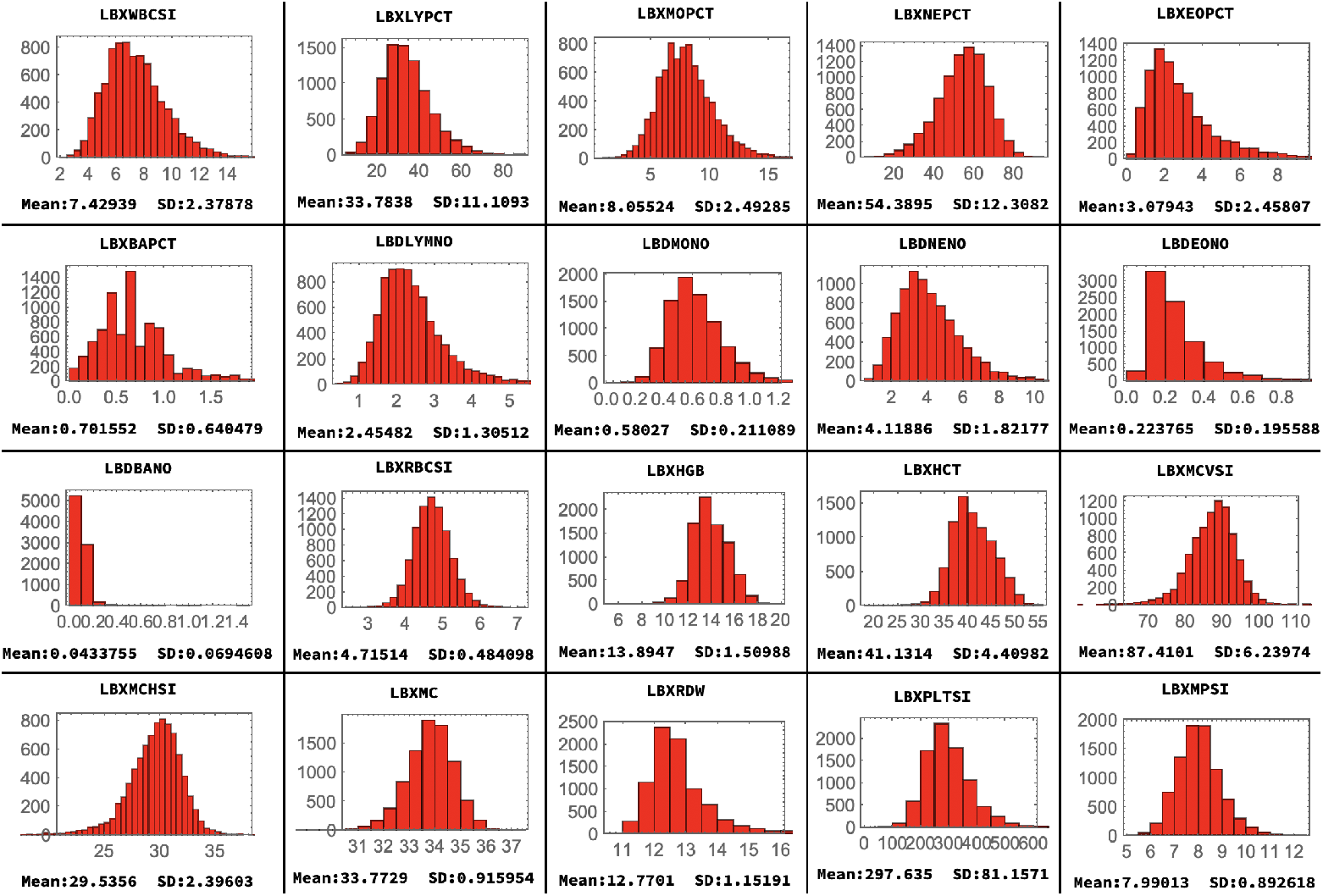
Values within ‘healthy’ reference ranges are heavily skewed and do not follow a normal or uniform distribution. Here, for illustration, are the aggregated distributions of the NHANES 2003 cohort per FBC analyte. This means that changes within what are considered normal ranges are different and thus more relevant across different analytes when medians and standard deviations are estimated empirically from the data. The list of analyte labels is in the Complementary Material.

## 5 Discussion and other aspects

### 5.1 Non-linear aspects and re-linearisation

We started with a simple version in which we assumed that there is no (common causal) influence between analytes during evaluation of the score. We then moved to a (theoretical) scenario where analyte values influence each other.

The scores explored are linear in the sense that no direct interaction between two analytes can affect the final result, unless both are off the normal range, and in proportion to the sum of the deviations.

One factor that’s key to capturing all the information encoded in a full blood test as performed by a senior medical professional is the interaction of the various analytes. As has been described so far, most aspects of the score are linear, except for the colour scheme rule that captures the gravity of 3 or more analytes outside the normal ranges. One way to keep the score linear while capturing non-linearities is by creating synthetic analytes. For example, the neutrophil-to-lymphocyte ratio relation is a key determinant of severity for Sepsis, and involves 2 analytes. The synthetic analyte that can be added is the ratio itself, thus replacing a non-linear rule that would make the score’s description eventually too convoluted to read with ease with a key marker as another analyte [5] and [6].

The introduction of weights as functions for each analyte also permits the incorporation of more medical expertise in the way in which the score is calculated.

## 6 Conclusion

We have introduced two risk-assessment scores and studied their behaviour against typical synthetic and empirical disease cases. Even when not designed to diagnose, but to flag out-of-boundary values, both scores were informative for disease based on out-of-range values removed from the median and boundaries of reference values, using as a metric the standard deviation as learned from empirical cases.

## Complementary Material

NHS healthy reference population values used in the calculation of the score tests are in Table 5.

List of label headers in the NHANES database for blood related analytes:

- LBXWBCSI White blood cell count (1000 cells/uL)
- LBXLYPCT Lymphocyte percent (%)
- LBXMOPCT Monocyte percent (%)
- LBXNEPCT Segmented neutrophils percent (%)
- LBXEOPCT Eosinophils percent (%)
- LBXBAPCT Basophils percent (%)
- LBDLYMNO Lymphocyte number
- LBDMONO Monocyte number
- LBDNENO Segmented neutrophils number
- LBDEONO Eosinophils number
- LBDBANO Basophils number
- LBXRBCSI Red blood cell count (million cells/uL)
- LBXHGB Haemoglobin (g/dL)
- LBXHCT Haematocrit (%)
- LBXMCVSI Mean cell volume (fL)
- LBXMCHSI Mean cell haemoglobin (pg)
- LBXMC MCHC (g/dL)
- LBXRDW Red cell distribution width (%)
- LBXPLTSI Platelet count SI (1000 cells/uL)
- LBXMPSI Mean platelet volume (fL)

